# Collective border cell migration requires the zinc transporter Catsup to limit endoplasmic reticulum stress

**DOI:** 10.1101/2021.04.04.438395

**Authors:** Xiaoran Guo, Wei Dai, Denise Montell

## Abstract

Collective cell migration is critical for normal development, wound healing, and in tumor progression and metastasis. Border cells in the Drosophila ovary provide a genetically tractable model to identify molecular mechanisms that drive this important cell behavior. In an unbiased screen for defects in border cell migration in mosaic clones, we identified a mutation in the *catsup* gene. Catsup, the Drosophila ortholog of Zip7, is a large, multifunctional, transmembrane protein of the endoplasmic reticulum (ER), which has been reported to negatively regulate catecholamine biosynthesis, to regulate Notch signaling, to function as a zinc transporter, and to limit ER stress. Here we report that *catsup* knockdown caused ER stress in border cells and that ectopic induction of ER stress was sufficient to block migration. Notch and EGFR trafficking were also disrupted. Wild type Catsup rescued the migration defect but point mutations known to disrupt the zinc ion transport of Zip7 did not. We conclude that migrating cells are particularly susceptible to defects in zinc transport and ER homeostasis.

## Introduction

Collective cell migration has emerged as a key driver of normal organ development, wound repair, and tumor metastasis^1–4^. Border cell migration in the Drosophila ovary provides a powerful in vivo model of collective cell migration that is amenable to unbiased genetic screening. Drosophila ovaries are composed of ovarioles, which are strings of egg chambers progressing through 14 stages of development to mature eggs (Fig.1A). Each egg chamber is composed of 16 germ cells including 15 nurse cells and one oocyte, which are surrounded by epithelial follicle cells. During stage 9 (Fig. 1B and C), 4-8 border cells are specified at the anterior end of the egg chamber, delaminate from the follicular epithelium, and migrate posteriorly, reaching the anterior border of the oocyte by stage 10.

**Fig. 1.**
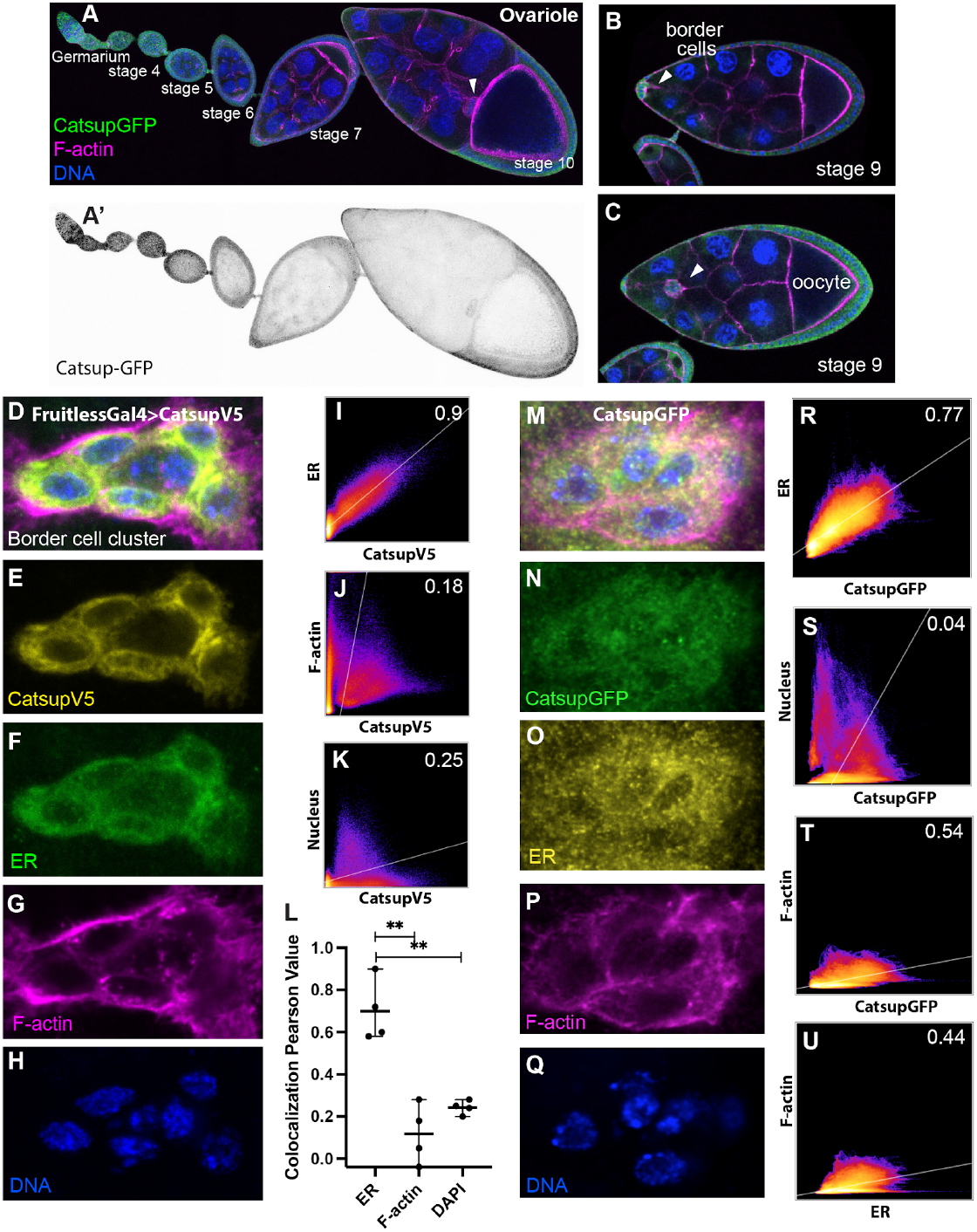
Catsup expression pattern in ovarioles. (A) An ovariole from germarium to developmental stage 10, where border cells (white arrowhead) arrive at the anterior border of the oocyte. (A’) An endogenously tagged Catsup::GFP shows the expression pattern of Catsup. (B) An early stage 9 egg chamber as border cells (arrowhead) initiate migration. (C) A mid stage 9 egg chamber with border cells en route to the oocyte. (D-H) High magnification of a border cell cluster showing the localization of overexpressed CatsupV5 (yellow), anti-PDI staining for ER (green), phalloidin staining for F-actin (magenta), Hoechst staining for DNA (blue). (I-K) 2-dimensional intensity histograms for two selected channels showing colocalization of CatsupV5 relative to ER, F-actin and nuclei. The colocalization regression Pearson’s coefficient displayed in the upper right corner. (L) CatsupV5 is highly colocalized with ER rather than F-actin or nuclei, shown by quantification of the Pearson’s coefficients from 4 border cells. ** P value < 0.01. (M-Q) High magnification of a border cell cluster showing the localization of endogenously tagged Catsup::GFP, ER (PDI), F-actin (Phalloidin) and nuclei (DNA). (R-U) 2-dimensional intensities histograms for two selected channels showing the colocalization and the Pearson’s coefficient of CatsupV5 relative to ER, nuclei, F-actin, and of ER relative to F-actin.

Genetic screens have yielded insights into the molecular mechanisms that specify which of the ~850 follicle cells acquire the ability to migrate^5,6^, the developmental timing of the migration^7,8^, collective direction sensing, and cytoskeletal dynamics^9–19^. While much is understood, insights continue to emerge from border cell studies^20–34^. The gene *catsup* was identified both in a large-scale, ethyl methanesulfonate-induced mutagenesis screen for border cell migration defects in mosaic clones^35^ and in a whole-genome gene expression profile^36^.

The name *catsup* is an abbreviation of “catecholamines up”, loss of which increases synthesis of aromatic amines including neurotransmitters such as epinephrine and dopamine^37^. Catsup is required for Drosophila tracheal morphogenesis, and in this context, it inhibits the Drosophila homolog of tyrosine hydroxylase (Ple) to limit dopamine synthesis^38^. In contrast, in wing imaginal disc cells, Catsup facilitates proper trafficking of Notch and EGFR^39^. Its mechanism of action in border cells is unknown.

Catsup shares a 62% similarity and 53% identity with its mammalian homolog ZIP7 (also known as SLC39A7 or HKE4)^39^, a member of one of the two major families of zinc transporters^40^. ZIP7 is located within intracellular membranes including ER where ZIP7 transports Zn^2+^ to the cytosol^41^. Zinc is a necessary trace element vital for many proteins to function, and zinc homeostasis requires 24 zinc transporters in humans, 14 of which are ZIPs^42^. Zip7 is a conserved protein found in the ER and Golgi in organisms as diverse as yeast, plants and animals^43–48^. In animal cells, loss of ZIP7 can lead to ER stress and in some cases cell death. Furthermore, increased ZIP7 expression is positively correlated with cancer cell proliferation, growth, invasion, and metastasis^49^ of breast^50,51^, cervical^52^ and colorectal cancer^53^.

In the mammalian intestine, loss of ZIP7 causes an increase in ER stress and loss of stem cells^54^. Similarly, catsup loss-of-function causes ER stress in fly wing imaginal discs^39^. Thus, Catsup and ZIP7 are multifunctional proteins. However, the relationships between ER stress, zinc transport, and cell motility remain to be clarified.

## Results

Using an endogenously-tagged Catsup::GFP fusion, we found that Catsup is expressed throughout oogenesis, including in all follicle cells (Fig. 1A-C). Mammalian ZIP7 localizes predominantly to the ER^41^, so we investigated the subcellular localization of Catsup. Both overexpressed, tagged CatsupV5 (Fig. 1D-L) and endogenously tagged Catsup::GFP (Fig. 1M-U) significantly co-localized with the ER resident enzyme protein disulfide isomerase (PDI), but not with DNA or F-actin. Expressing a Catsup RNAi line in border cells using c306Gal4 (Fig. 2A) and FruitlessGal4 led to incomplete migration in 80% of stage 10 egg chambers examined (Fig.1B and C), a defect that was rescued by UAS-CatsupV5 (Fig. 1C). Using the FLP-FRT system, we generated clones of *catsup* mutant cells in genetically mosaic egg chambers. Compared to control clones (Fig. 2D-D”), border cell clusters containing cells homozygous mutant for catsup exhibited migration defects (Fig. 2E-E”), the severity of which was proportional to the percentage of mutant cells per cluster (Fig. 2F). In addition, in clusters containing both heterozygous and homozygous mutant cells, homozygous mutant cells tended to occupy rear positions (Fig. 2G-G’ and E), which is typical of mutations in genes required for motility^55^.

**Fig. 2.**
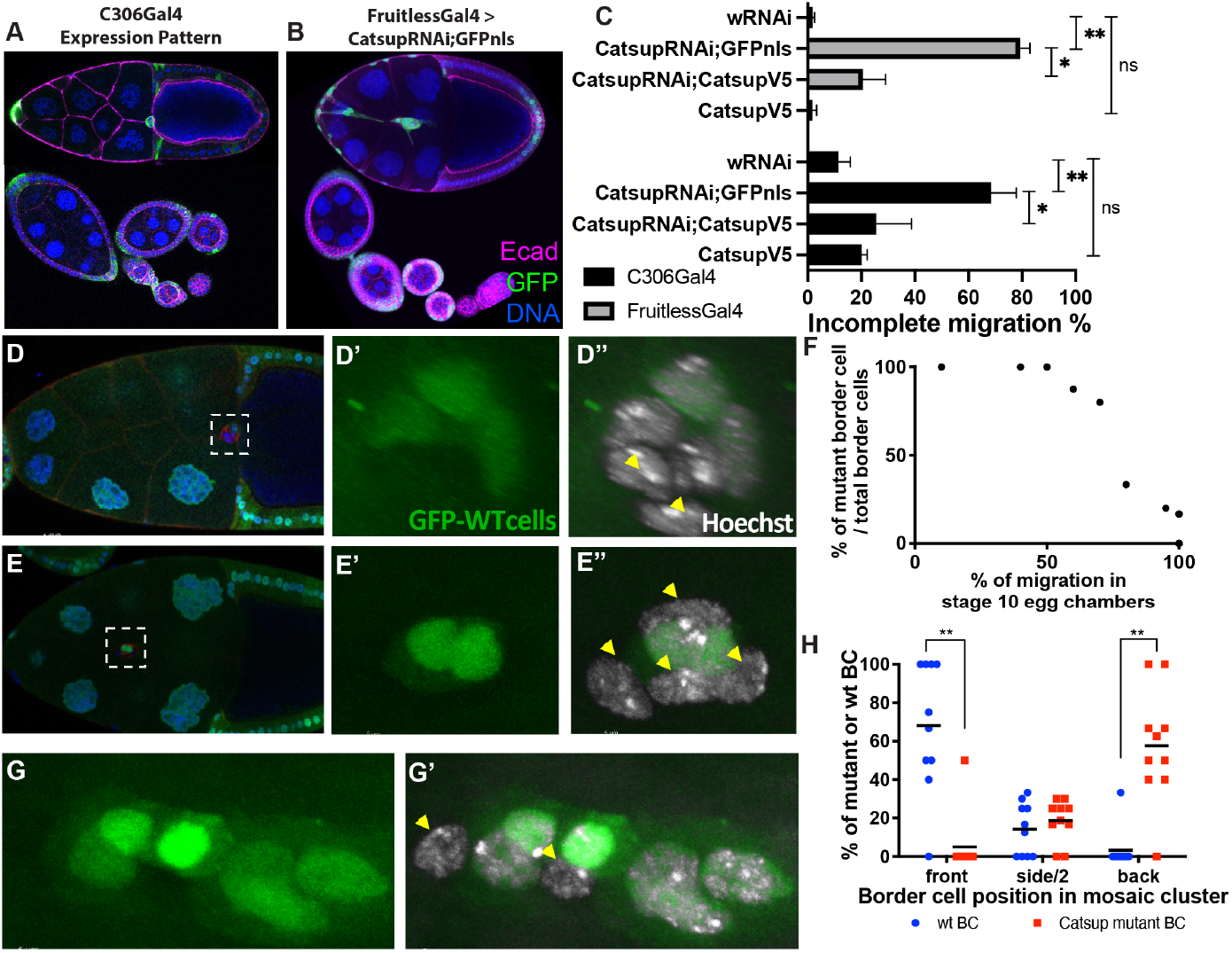
Catsup is required for border cell migration. (A) C306Gal4 drives UAS-transgene expression in anterior and posterior follicle cells throughout development, including in border cells. (B) An example of incomplete migration caused by CatsupRNAi, expressed with fruitlessGal4 in anterior and posterior follicle cells including border cells, but not polar cells. (C) Quantification of the incomplete migration at stage 10 in C306Gal4 (black bars) and FruitlessGal4 (grey bars) driving the negative control (wRNAi) or UAS-CatsupRNAi;UAS-GFPnls. UAS-CatsupV5 significantly rescues the defect. CatsupV5 alone does not cause more incomplete migration than controls. (D) Control clones in which homozygous yellow mutant cells are GFP-, and the wild type cells are GFP+ green cells. (E) GFP-negative cells are homozygous catsup mutant border cells. (F) The migration distance expressed as a percentage of the migration path for mosaic border cell clusters. Migration impairment is related to the proportion of homozygous mutant cells in the cluster. (G) Two mutant border cells are lagging behind the migration cluster. (H) Quantification shows mutant border cells locating in the opposite direction of migration. Yellow arrowheads point at mutant border cells.

One known function of Catsup is direct binding to and inhibition of the tyrosine hydroxylase Ple, which is the rate-limiting enzyme in catecholamine synthesis^56^. Ple and Catsup are both expressed in embryonic tracheal cells, where they contribute to achieving proper dopamine levels, which regulate Breathless (fibroblast growth factor receptor) endocytosis and signaling^38^. To test whether inhibition of Ple by Catsup was critical for border cells to migrate, we used an antibody to assess Ple expression in wild-type egg chambers. In contrast to tracheal cells, we detected no Ple protein in wild-type egg chambers (Fig. 3A). The antibody was effective because we could detect Ple ectopically expressed using c306Gal4 (Fig. 3B), as well as endogenous expression of Ple in neurons in the adult brain^57^ (Fig. 3C). Furthermore, Ple overexpression in border cells caused no migration defect (Fig. 3B). Therefore, it is unlikely that negative regulation of Ple activity is the key function of Catsup in border cells, suggesting that the function of Catsup in border cell migration is likely distinct from its role in tracheal development.

**Fig. 3.**
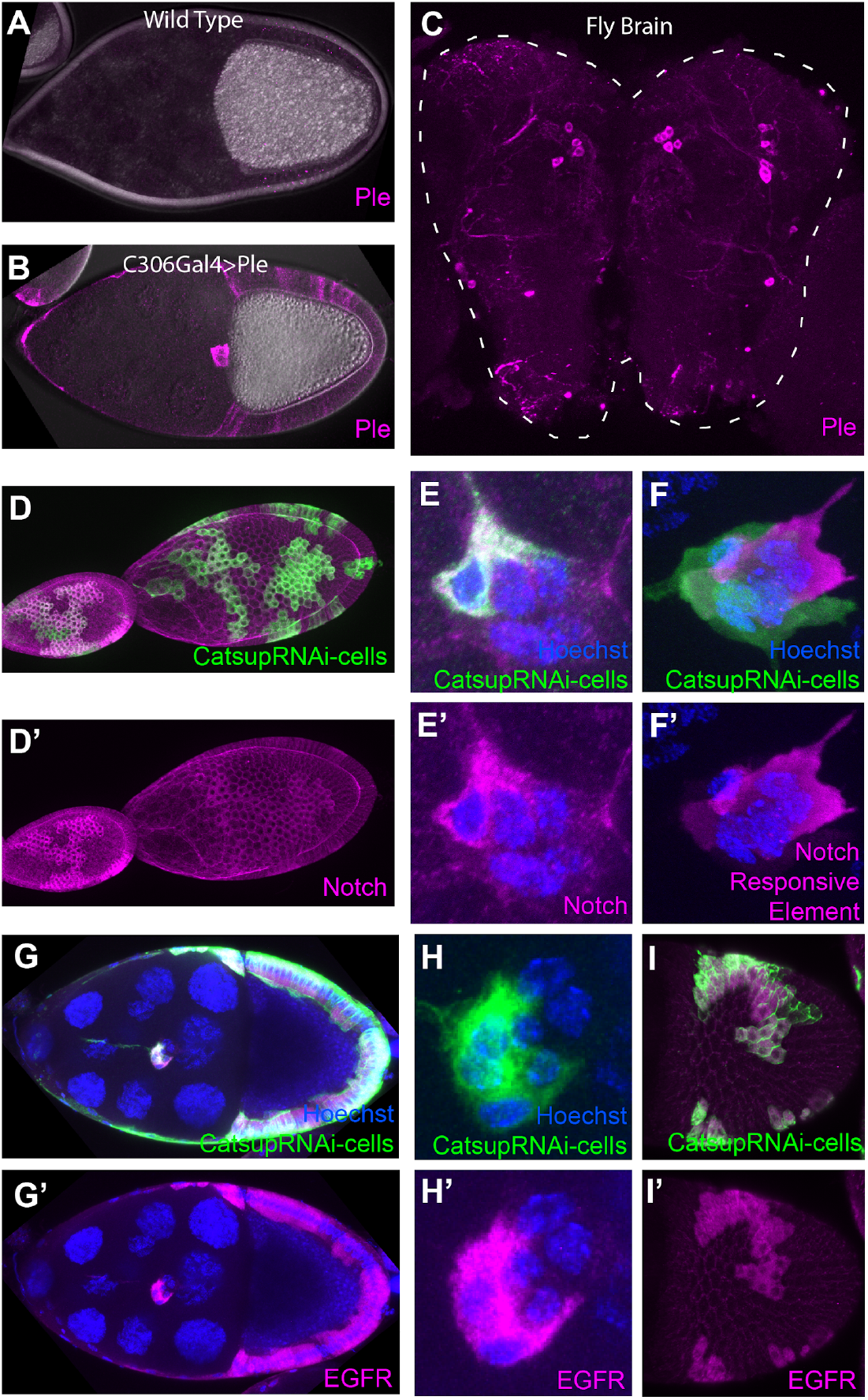
Catsup loss of function impairs Notch signaling. (A, B) Differential interference contrast images of stage 10 egg chambers stained for Ple (magenta). (A) A wild type egg chamber. (B) a stage 10 egg chamber expressing UAS-Ple with c306Gal4. (C) A dissected fly brain stained for endogenous Ple: dopaminergic neurons are Ple-positive. (D-E’) CatsupRNAi-expressing clones (GFP-positive, green) accumulate intracellular Notch protein in epithelial follicle cells (D, D’) and border cells (E, E’) relative to neighboring wild type cells. (F-F’) CatsupRNAi-expressing border cells (GFP-positive, green) show decreased Notch signaling shown by the Notch-responsive-element driving RFP (magenta)relative to neighboring wild type cells. (G-I) Accumulation of EGFR (magenta) in CatsupRNAi-expressing epithelial follicle cells (G, G’ and I, I’) and border cells H, H’).

In wing imaginal discs, Notch and EGFR signaling are disrupted by Catsup loss of function, and both of these pathways are required for border cell migration. Notch signaling facilitates initiation of border cell migration, specifically detachment from the anterior^58^, while EGFR is required for border cells to take a dorsal turn near the end of their migration to the oocyte^16,23^. We found abnormal intracellular accumulation of Notch in epithelial cells generally (Fig. 3D, D’) and border cells specifically (Fig. 3E, E’) upon Catsup knockdown (Fig. 3D and E). Cells lacking Catsup also exhibited defective Notch signaling, detected by the Notch responsive element reporter^59^ (Fig. 3E, E’). As in imaginal discs, EGFR also accumulated abnormally (Fig. 3G-I’). Since Notch signaling is essential for border cell migration and expression of constitutively active Notch (the Notch intracellular domain, NICD), which does not require intracellular trafficking or processing, rescues impaired Notch signaling in border cells^58^, we asked whether NICD expression might rescue Catsup knockdown. However, neither NICD expression nor overexpression of the Notch specific chaperone O-fucosyltransferase-1^60^ was sufficient to rescue Catsup RNAi (Fig. S1). This suggests that multiple Catsup functions are essential for border cell migration.

Since suppression of ER stress is a conserved function of Catsup and ZIP7, we used the ER stress marker XBP1-EGFP^61^ to compare heterozygous and homozygous catsup mutant cells in mosaic clusters (Fig. 4A). We found high levels of Xbp1 protein in homozygous cells compared to heterozygous border cells. Cells with elevated Xbp1 also exhibited reduced expression of Eyes Absent (Eya) (Fig. 4A), a nuclear protein that is required to repress polar cell fate and maintain border cell identity^6^. By contrast, the nuclear protein STAT, a transcription factor required for border cell fate specification^5^, was not decreased. Nuclear size was reduced by about half in mutant cells, possibly due to defective Notch signaling^62,63^ (Fig. 4B).

**Fig. 4.**
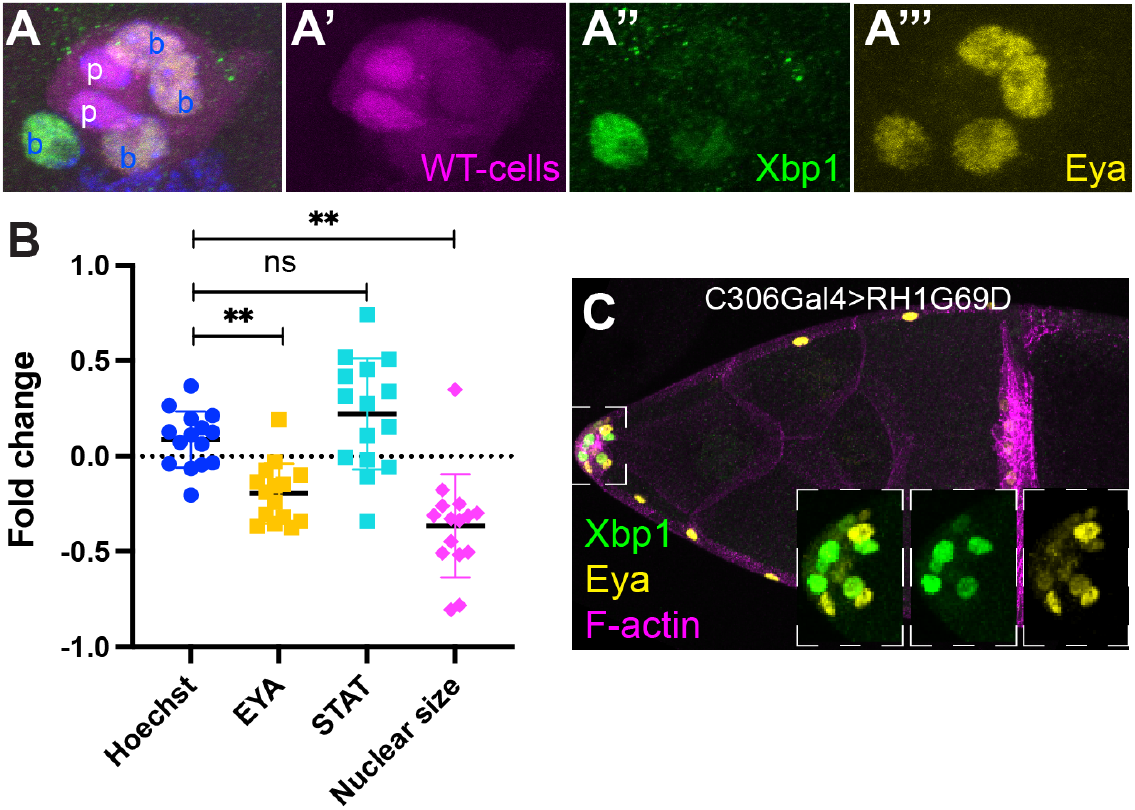
Catsup RNAi causes ER stress in border cells. (A) A mosaic border cell cluster composed of heterozygous and homozygous catsup mutant cells (RFP-positive cells, magenta, are catsup-/+; RFP-negative cells are catsup-/-). Wild type polar cells (p) express a higher level of RFP compared to wild type border cells (b). The homozygous catsup mutant border cell shows elevated expression of Xbp1::EGFP (green), a marker for ER stress. Eya (yellow) is a cell fate and differentiation marker for border cells that is absent from polar cells and is reduced in the homozygous catsup mutant border cell. (B) Quantification of fluorescence intensity per unit area for DNA (Hoechst), Eya, and STAT and of nuclear size expressed as a fold change between homozygous mutant and heterozygous cells. (C) Mosaic overexpression of an misfolded mutant rhodopsin protein RH1^G69D^ in border cells caused ER stress (Xbp1::EGFP in green), decreased Eya (yellow), and blocked border cell migration.

To test whether ER stress impairs migration, we genetically induced ER stress by overexpressing a misfolded rhodopsin protein RH1^G69D 64^. As expected, RH1^G69D^ induced Xbp1 expression in border cells (Fig. 4C). It also blocked migration (Fig. 4C), showing that high levels of ER stress are sufficient to inhibit border cell migration. In Xbp1-positive, RH1^G69D^-expressing cells, Eya expression was again reduced (Fig. 4C, insets), suggesting that ER stress, rather than another function of Catsup, was likely responsible for reduced Eya protein levels.

ZIP7 transports zinc from the ER into the cytosol^65^. To test whether the zinc transporter activity of Catsup was likely required for border cell migration, we designed point mutations that alter amino acid residues that are conserved between Catsup, ZIP7 and a more distant family member from Arabidopsis IRT1, and that are required for zinc transport (Fig. 5B). Histidine is an amino acid that coordinates zinc^66^, and it appears in the highly conserved HELP domain and the CHEXPHEXGD motif that are important for Zinc transport^40^. The mutation Catsup^H315A^ replaces a conserved histidine within the HELP domain with alanine. Catsup^H344A^ changes a histidine within the CHEXPHEXGD motif to alanine. We made transgenic flies expressing the mutants under Gal4 control and included a V5 tag so that we could monitor protein expression and localization. As a control, we made UAS-Catsup^G178D^, the point mutation present in the mutant line (Catsup^68E2^) isolated from the screen for mutations that cause border cell migration defects in mosaic clones^35^. We then co-expressed each of the mutant lines with CatsupRNAi and quantified border cell migration (Fig. 5C). UAS-Catsup^G178D^ provided no significant rescue compared to UAS-GFP-nls, though the mutation is likely a hypomorph and may have provided slight rescue that did not reach statistical significance (P=0.1). Castup^H344A^ failed to rescue, as did expression of UAS-Catsup^H315A^, which though not statistically significantly different, might have caused an even more severe migration defect, possibly due to a dominant-negative effect (Fig. 5C). The point mutations did not destabilize the proteins or alter their localization (Fig. 5D, E), therefore the lack of rescue was likely a consequence of impaired activity rather than impaired expression.

**Fig. 5.**
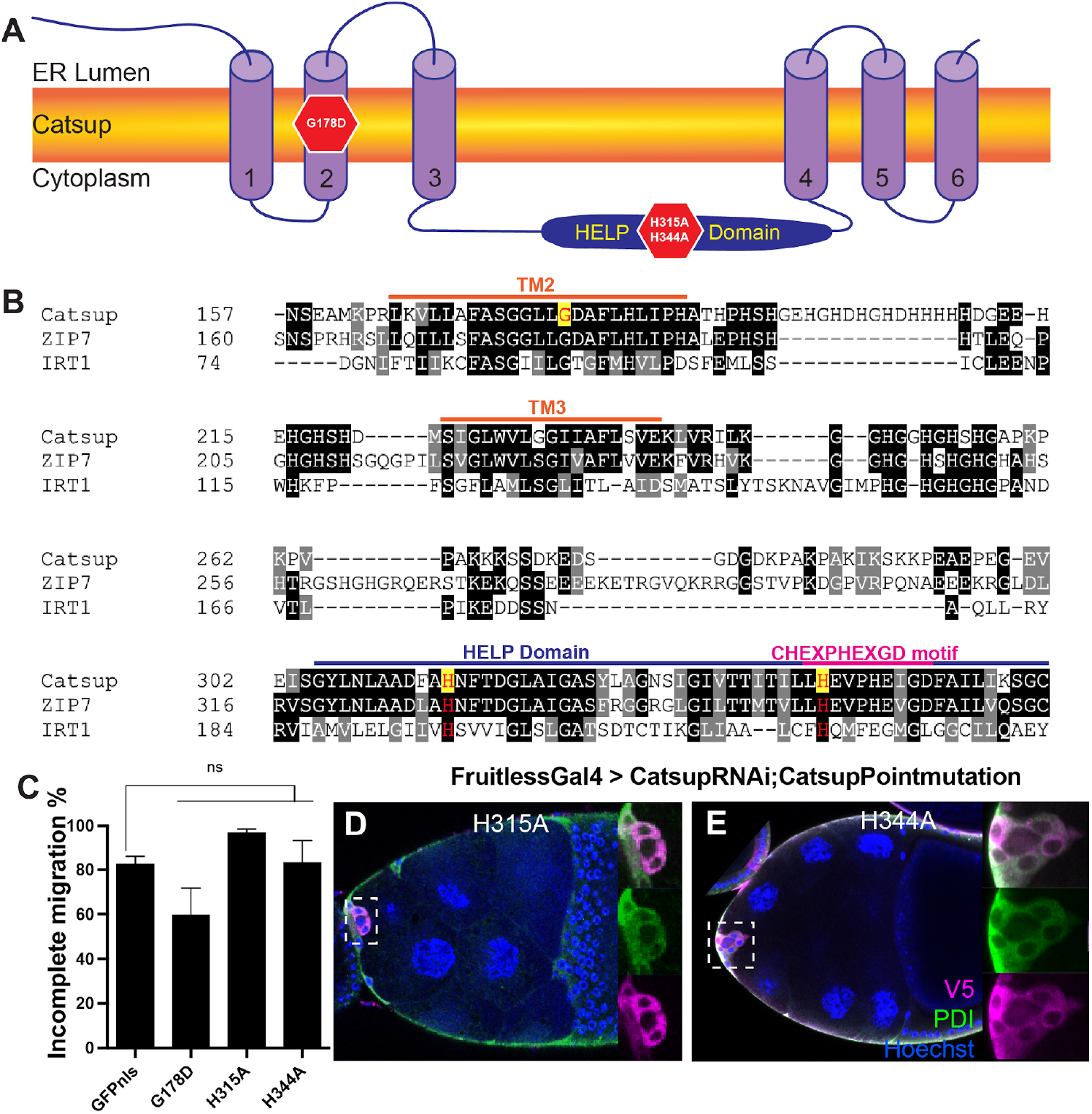
Point mutations reveal likely requirement for Zinc transport in cell motility. (A) Schematic representation of transmembrane domains and topology for ZIP family ion transporters (B) Sequence alignment of Catsup, ZIP7, and the arabidopsis protein IRT1. Identical amino acids are shaded in black, similar amino acids in gray. Amino acid substitutions generated in this study are highlighted in yellow with red text. (C) Quantification of incomplete migration at stage 10 in egg chambers expressing catsupRNAi together with the indicated mutant forms. (D-E) Expression and co-localization of CatsupH315A and CatsupH344A with the ER marker PDI in border cells.

## Discussion

In this study we explored the roles of the multifunctional protein Catsup in border cell migration. Catsup is a conserved protein that goes by names including ZRT1 in yeast, IRT1 in plants, SLC39a7/Zip7/Ke4 in mammals. Despite the name Catsup (Catecholamines up), the most conserved features include subcellular localization to intracellular membranes including ER and Golgi, and bivalent cation transport.

In Drosophila, Catsup has been implicated in direct binding and inhibition of tyrosine hydroxylase (TH/Ple), and thus in limiting dopamine production. This is important in the nervous system where as a neurotransmitter, dopamine levels must be tightly regulated. Somewhat surprisingly, the negative regulation of dopamine by Catsup is also important in tracheal development. Hsouna et al showed that Catsup and Ple are both required to achieve the appropriate level of dopamine, which regulates internalization of the Breathless (Btl) receptor tyrosine kinase. Excessive dopamine in *catsup* mutants leads to excessive endocytosis and thus downregulation of Btl, which inhibits tracheal cell migration. Ple mutations on the other hand result in reduced dopamine and Btl endocytosis, excess Btl signaling and therefore ectopic branching. Border cells also rely on chemotaxis via receptor tyrosine kinase signaling, so our first hypothesis was that the mechanism would be similar in tracheal and border cells. However, Ple is not detectable in border cells, so it is unlikely that negative regulation of dopamine synthesis is the primary function of Catsup in border cells.

In Drosophila wing imaginal discs, Catsup was found to regulate trafficking of Notch and EGFR. Although the biological effects of Notch and EGFR are different in imaginal disc cells compared to border cells, both pathways are required in both cell types, and our results support a general role for Catsup in Notch and EGFR trafficking. Perhaps as a consequence of defective trafficking, Catsup knockdown also induced ER stress in border cells, supporting that this is a general and conserved function^54^. Moreover, we show for the first time that ER stress is sufficient to block migration and thereby regulate the abundance of the nuclear protein Eya, whereas STAT is unaffected. It is not immediately obvious how ER stress affects Eya abundance. Eya is an unusual protein in that it possesses protein phosphatase activity^67,68^, at least in vitro, and serves as a transcriptional activator^69^. Together the results demonstrate multiple essential functions for Catsup in border cells.

The point mutations designed to disrupt zinc transport based on equivalent mutations in ZIP7 indicated that border cell migration requires not only expression and ER localization of Catsup but also its ability to transport zinc. Given the requirement for ZIP7 in cancer cell motility. its over-expression in numerous cancers, and its correlation with disease progression, invasion, and metastasis, the border cell system offers an excellent model for deciphering the key effects of this multifunctional protein on collective cell motility in vivo.

## Materials and Methods

### Drosophila genetics

Catsup mutant fly was generated by ethyl methanesulfonate (EMS) mutagen. The mutation results in Glycine(G) to Aspartic acid(D) replacement at the 178th amino acid. FLP/FRT system was used to generate the CatsupG178D homozygous mutant clones by combining FRT40A-CatsupG178D with hsFLP12,yw;ubi:GFPnls, FRT40A or hsFLP12,yw;ubi:RFPnls, FRT40A/(CyO). Catsup knock down experiment uses border cell specific C306Gal4 and FruitlessGal4 to drive UASCatsupRNAi that is from VDRC 100095 P{KK103630}VIE-260B. Wild type rescue w[*]; sna[Sco]/CyO; P{w[+mC]=UAS-Catsup.V5}6 Bloomington 63229. Transgenic drosophila stocks used UASwRNAi/Cyo is a lab stock, UASPleRNAi Bloomington 25796 y[1] v[1]; P{y[+t7.7] v[+t1.8]=TRiP.JF01813}attP2, UASPle is Bloomington 37539 w[*]; P{w[+mC]=UAS-ple.T}331f2, O-fucosyltransferase1 Bloomington 9376 P{UAS-O-fut1.O}11.1, UAS-Notch.Intracellular.Domain on third chromosome is a gift from Artavanis-Tsakonas Lab^70^, ER stress marker UASXbp1-EGFP.HG Bloomington 60731 w[*]; P{w[+mC]=UAS-Xbp1.EGFP.HG}3, UASHSC70-3 Bloomington 5843 w[126]; P{w[+mC]=UAS-Hsc70-3.WT}B. Catsup point mutations were cloned into vector pUASt-attb and injected to attp2 flies y1 w67c23; P{CaryP}attP2 by BestGene Inc.

### Design of UAS-RNAi-resistant Catsup point mutations

When generating UAS-Catsup-point-mutations, we designed the construct so it can not be targeted by the CatsupRNAi sequences. The region that is targeted by RNAi changed with the redundant codon for the same amino acids.

### Immunostaining and confocal imaging

Female flies were fattened with yeast for 2 days at 29°C. Egg chambers are dissected from ovaries of female fly bodies in Schneider’s medium with 10% FBS (pH=6.85-6.95) as described previously^71^. Freshly dissected egg chambers are fixed in 4% paraformaldehyde and then incubated overnight in 1xPBS with 0.4% triton with primary antibody to stain for ER mouse PDI (1:200) ADI-SPA-891-D Enzo Life Sciences, Inc., chicken GFP (1:200) ab13970 Abcam plc., Ple (anti-TH) antibody is a gift from Craig Montell lab, mouse anti-Notch intracellular domain (1:100) C17.9C6 DSHB, rat Ecadherin antibody DCAD2 (1:50) DSHB, V5 Tag Monoclonal Antibody-Alexa Fluor 555 (2F11F7) Invitrogen, mouse anti-dEGFR (1:2000) E2906 Sigma Aldrich. O-fut1 antibody was used to confirm O-fut1 overexpression, and it is a gift from Kenneth D. Irvine lab^72^. Secondary antibodies were incubated for 2 hours, together with Hoechst stains for nuclei, and Phalloidin stains for F-actin. Mouse anti-PDI and mouse anti-V5-555 co-staining was done by first stain with PDI primary and secondary, after through wash out, apply anti-V5-555 for overnight then wash out. Immunostained samples are mounted in VECTASHIELD Mounting Medium from Vector Laboratories. Zeiss LSM780 and LSM800 confocal microscopes were used to acquire images. Images were visualised by FIJI, rotated and cropped for presentation.

### Sequence alignment

Catsup and ZIP7 amino acid sequences were acquired from NCBI in a FASTA format. The files were input into T-coffee http://tcoffee.crg.cat/apps/tcoffee/do:regular to generate multiple sequence alignment. The output was fed into Boxshade http://www.ch.embnet.org/software/BOX_form.html to generate the sequence alignment with black and grey shades to show conserved sequence region.

## End Matter

### Author Contributions and Notes

X.G., W.D., and D.J.M. designed experiments and prepared the manuscript. X.G., and W.D., performed experiments.

The authors declare no conflict of interest.

## Acknowledgments

We thank Kristin Mercier, Marc Anthony Pastor and Yijing Li for their technical assistance. Thanks to all the colleagues in Denise Montell lab for their useful suggestions and kind helps. We thank the labs that generously shared the reagents with us.

## References

1. Mishra, A. K., Campanale, J. P., Mondo, J. A. & Montell, D. J. Cell interactions in collective cell migration. Development 146, (2019).

2. Friedl, P. & Mayor, R. Tuning Collective Cell Migration by Cell-Cell Junction Regulation. Cold Spring Harb. Perspect. Biol. 9, (2017).

3. Friedl, P. & Gilmour, D. Collective cell migration in morphogenesis, regeneration and cancer. Nat. Rev. Mol. Cell Biol. 10, 445–457 (2009).

4. Friedl, P., Locker, J., Sahai, E. & Segall, J. E. Classifying collective cancer cell invasion. Nat. Cell Biol. 14, 777–783 (2012).

5. Silver, D. L. & Montell, D. J. Paracrine signaling through the JAK/STAT pathway activates invasive behavior of ovarian epithelial cells in Drosophila. Cell 107, 831–841 (2001).

6. Bai, J. & Montell, D. Eyes absent, a key repressor of polar cell fate during Drosophila oogenesis. Development 129, 5377–5388 (2002).

7. Jang, A. C.-C., Chang, Y.-C., Bai, J. & Montell, D. Border-cell migration requires integration of spatial and temporal signals by the BTB protein Abrupt. Nat. Cell Biol. 11, 569–579 (2009).

8. Bai, J., Uehara, Y. & Montell, D. J. Regulation of invasive cell behavior by taiman, a Drosophila protein related to AIB1, a steroid receptor coactivator amplified in breast cancer. Cell 103, 1047–1058 (2000).

9. Murphy, A. M. & Montell, D. J. Cell type-specific roles for Cdc42, Rac, and RhoL in Drosophila oogenesis. J. Cell Biol. 133, 617–630 (1996).

10. Kim, J. H. et al. Psidin, a conserved protein that regulates protrusion dynamics and cell migration. Genes Dev. 25, 730–741 (2011).

11. McDonald, J. A., Pinheiro, E. M. & Montell, D. J. PVF1, a PDGF/VEGF homolog, is sufficient to guide border cells and interacts genetically with Taiman. Development 130, 3469–3478 (2003).

12. Lee, T., Feig, L. & Montell, D. J. Two distinct roles for Ras in a developmentally regulated cell migration. Development 122, 409–418 (1996).

13. Cai, D. et al. Mechanical feedback through E-cadherin promotes direction sensing during collective cell migration. Cell 157, 1146–1159 (2014).

14. McDonald, J. A., Khodyakova, A., Aranjuez, G., Dudley, C. & Montell, D. J. PAR-1 kinase regulates epithelial detachment and directional protrusion of migrating border cells. Curr. Biol. 18, 1659–1667 (2008).

15. Wang, X., He, L., Wu, Y. I., Hahn, K. M. & Montell, D. J. Light-mediated activation reveals a key role for Rac in collective guidance of cell movement in vivo. Nat. Cell Biol. 12, 591–597 (2010).

16. Duchek, P. & Rørth, P. Guidance of cell migration by EGF receptor signaling during Drosophila oogenesis. Science 291, 131–133 (2001).

17. Fulga, T. A. & Rørth, P. Invasive cell migration is initiated by guided growth of long cellular extensions. Nat. Cell Biol. 4, 715–719 (2002).

18. Ramel, D., Wang, X., Laflamme, C., Montell, D. J. & Emery, G. Rab11 regulates cell-cell communication during collective cell movements. Nat. Cell Biol. 15, 317–324 (2013).

19. Assaker, G., Ramel, D., Wculek, S. K., González-Gaitán, M. & Emery, G. Spatial restriction of receptor tyrosine kinase activity through a polarized endocytic cycle controls border cell migration. Proc Natl Acad Sci USA 107, 22558–22563 (2010).

20. Miao, G., Godt, D. & Montell, D. J. Integration of Migratory Cells into a New Site In Vivo Requires Channel-Independent Functions of Innexins on Microtubules. Dev. Cell 54, 501–515.e9 (2020).

21. Colombié, N. et al. Non-autonomous role of Cdc42 in cell-cell communication during collective migration. Dev. Biol. 423, 12–18 (2017).

22. Chen, Y. et al. Protein phosphatase 1 activity controls a balance between collective and single cell modes of migration. elife 9, (2020).

23. Dai, W. et al. Tissue topography steers migrating Drosophila border cells. BioRxiv (2020) doi:10.1101/2020.09.27.316117.

24. Laflamme, C. et al. Evi5 promotes collective cell migration through its Rab-GAP activity. J. Cell Biol. 198, 57–67 (2012).

25. Ogienko, A. A. et al. GAGA regulates border cell migration in drosophila. Int. J. Mol. Sci. 21, (2020).

26. Berez, A., Peercy, B. E. & Starz-Gaiano, M. Development and analysis of a quantitative mathematical model of bistability in the cross repression system between APT and SLBO within the JAK/STAT signaling pathway. Front. Physiol. 11, 803 (2020).

27. Wang, X. et al. Temporal Coordination of Collective Migration and Lumen Formation by Antagonism between Two Nuclear Receptors. iScience 23, 101335 (2020).

28. Wang, H., Guo, X., Wang, X., Wang, X. & Chen, J. Supracellular Actomyosin Mediates Cell-Cell Communication and Shapes Collective Migratory Morphology. iScience 23, 101204 (2020).

29. Fox, E. F., Lamb, M. C., Mellentine, S. Q. & Tootle, T. L. Prostaglandins regulate invasive, collective border cell migration. Mol. Biol. Cell 31, 1584–1594 (2020).

30. Plutoni, C. et al. Misshapen coordinates protrusion restriction and actomyosin contractility during collective cell migration. Nat. Commun. 10, 3940 (2019).

31. Zeledon, C., Sun, X., Plutoni, C. & Emery, G. The ArfGAP Drongo Promotes Actomyosin Contractility during Collective Cell Migration by Releasing Myosin Phosphatase from the Trailing Edge. Cell Rep. 28, 3238–3248.e3 (2019).

32. Lamb, M. C., Anliker, K. K. & Tootle, T. L. Fascin regulates protrusions and delamination to mediate invasive, collective cell migration in vivo. Dev. Dyn. 249, 961–982 (2020).

33. Ghiglione, C., Jouandin, P., Cérézo, D. & Noselli, S. The Drosophila insulin pathway controls Profilin expression and dynamic actin-rich protrusions during collective cell migration. Development 145, (2018).

34. Sharma, A. et al. Insulin signaling modulates border cell movement in Drosophila oogenesis. Development 145, (2018).

35. Liu, Y. & Montell, D. J. Identification of mutations that cause cell migration defects in mosaic clones. Development 126, 1869–1878 (1999).

36. Wang, X. et al. Analysis of cell migration using whole-genome expression profiling of migratory cells in the Drosophila ovary. Dev. Cell 10, 483–495 (2006).

37. Stathakis, D. G. et al. The catecholamines up (Catsup) protein of Drosophila melanogaster functions as a negative regulator of tyrosine hydroxylase activity. Genetics 153, 361–382 (1999).

38. Hsouna, A., Lawal, H. O., Izevbaye, I., Hsu, T. & O’Donnell, J. M. Drosophila dopamine synthesis pathway genes regulate tracheal morphogenesis. Dev. Biol. 308, 30–43 (2007).

39. Groth, C., Sasamura, T., Khanna, M. R., Whitley, M. & Fortini, M. E. Protein trafficking abnormalities in Drosophila tissues with impaired activity of the ZIP7 zinc transporter Catsup. Development 140, 3018–3027 (2013).

40. Kambe, T. Zinc Transport: Regulation. in Encyclopedia of inorganic and bioinorganic chemistry (ed. Scott, R. A.) 1–9 (John Wiley & Sons, Ltd, 2011). doi:10.1002/9781119951438.eibc2135.

41. Taylor, K. M., Morgan, H. E., Johnson, A. & Nicholson, R. I. Structurefunction analysis of HKE4, a member of the new LIV-1 subfamily of zinc transporters. Biochem. J. 377, 131–139 (2004).

42. Kambe, T., Tsuji, T., Hashimoto, A. & Itsumura, N. The physiological, biochemical, and molecular roles of zinc transporters in zinc homeostasis and metabolism. Physiol. Rev. 95, 749–784 (2015).

43. Nguyen, T. S. L., Kohno, K. & Kimata, Y. Zinc depletion activates the endoplasmic reticulum-stress sensor Ire1 via pleiotropic mechanisms. Biosci. Biotechnol. Biochem. 77, 1337–1339 (2013).

44. Tan, S. et al. Over-expression of the MxIRT1 gene increases iron and zinc content in rice seeds. Transgenic Res. 24, 109–122 (2015).

45. Zhang, P. et al. An uncleaved signal peptide directs the Malus xiaojinensis iron transporter protein Mx IRT1 into the ER for the PM secretory pathway. Int. J. Mol. Sci. 15, 20413–20433 (2014).

46. Adulcikas, J., Norouzi, S., Bretag, L., Sohal, S. S. & Myers, S. The zinc transporter SLC39A7 (ZIP7) harbours a highly-conserved histidine-rich N-terminal region that potentially contributes to zinc homeostasis in the endoplasmic reticulum. Comput. Biol. Med. 100, 196–202 (2018).

47. Tuncay, E. et al. Hyperglycemia-Induced Changes in ZIP7 and ZnT7 Expression Cause Zn2+ Release From the Sarco(endo)plasmic Reticulum and Mediate ER Stress in the Heart. Diabetes 66, 1346–1358 (2017).

48. Fauster, A. et al. Systematic genetic mapping of necroptosis identifies SLC39A7 as modulator of death receptor trafficking. Cell Death Differ. 26, 1138–1155 (2019).

49. Hogstrand, C., Kille, P., Nicholson, R. I. & Taylor, K. M. Zinc transporters and cancer: a potential role for ZIP7 as a hub for tyrosine kinase activation. Trends Mol. Med. 15, 101–111 (2009).

50. Taylor, K. M. et al. ZIP7-mediated intracellular zinc transport contributes to aberrant growth factor signaling in antihormone-resistant breast cancer Cells. Endocrinology 149, 4912–4920 (2008).

51. Ziliotto, S. et al. Activated zinc transporter ZIP7 as an indicator of antihormone resistance in breast cancer. Metallomics 11, 1579–1592 (2019).

52. Wei, Y., Dong, J., Li, F., Wei, Z. & Tian, Y. Knockdown of SLC39A7 suppresses cell proliferation, migration and invasion in cervical cancer. EXCLI J. 16, 1165–1176 (2017).

53. Sheng, N. et al. Knockdown of SLC39A7 inhibits cell growth and induces apoptosis in human colorectal cancer cells. Acta Biochim Biophys Sin (Shanghai) 49, 926–934 (2017).

54. Ohashi, W. et al. Zinc Transporter SLC39A7/ZIP7 Promotes Intestinal Epithelial Self-Renewal by Resolving ER Stress. PLoS Genet. 12, e1006349 (2016).

55. Haas, P. & Gilmour, D. Chemokine signaling mediates self-organizing tissue migration in the zebrafish lateral line. Dev. Cell 10, 673–680 (2006).

56. Wang, Z. et al. Catecholamines up integrates dopamine synthesis and synaptic trafficking. J. Neurochem. 119, 1294–1305 (2011).

57. Navarro, J. A. et al. Analysis of dopaminergic neuronal dysfunction in genetic and toxin-induced models of Parkinson’s disease in Drosophila. J. Neurochem. 131, 369–382 (2014).

58. Wang, X., Adam, J. C. & Montell, D. Spatially localized Kuzbanian required for specific activation of Notch during border cell migration. Dev. Biol. 301, 532–540 (2007).

59. Terriente-Felix, A. et al. Notch cooperates with Lozenge/Runx to lock haemocytes into a differentiation programme. Development 140, 926–937 (2013).

60. Okajima, T., Xu, A., Lei, L. & Irvine, K. D. Chaperone activity of protein O-fucosyltransferase 1 promotes notch receptor folding. Science 307, 1599–1603 (2005).

61. Sone, M., Zeng, X., Larese, J. & Ryoo, H. D. A modified UPR stress sensing system reveals a novel tissue distribution of IRE1/XBP1 activity during normal Drosophila development. Cell Stress Chaperones 18, 307–319 (2013).

62. Sun, J., Smith, L., Armento, A. & Deng, W.-M. Regulation of the endocycle/gene amplification switch by Notch and ecdysone signaling. J. Cell Biol. 182, 885–896 (2008).

63. Deng, W. M., Althauser, C. & Ruohola-Baker, H. Notch-Delta signaling induces a transition from mitotic cell cycle to endocycle in Drosophila follicle cells. Development 128, 4737–4746 (2001).

64. Chow, C. Y., Avila, F. W., Clark, A. G. & Wolfner, M. F. Induction of excessive endoplasmic reticulum stress in the Drosophila male accessory gland results in infertility. PLoS ONE 10, e0119386 (2015).

65. Taylor, K. M., Hiscox, S., Nicholson, R. I., Hogstrand, C. & Kille, P. Protein kinase CK2 triggers cytosolic zinc signaling pathways by phosphorylation of zinc channel ZIP7. Sci. Signal. 5, ra11 (2012).

66. Laitaoja, M., Valjakka, J. & Jänis, J. Zinc coordination spheres in protein structures. Inorg. Chem. 52, 10983–10991 (2013).

67. Rayapureddi, J. P. et al. Eyes absent represents a class of protein tyrosine phosphatases. Nature 426, 295–298 (2003).

68. Tootle, T. L. et al. The transcription factor Eyes absent is a protein tyrosine phosphatase. Nature 426, 299–302 (2003).

69. Li, X. et al. Eya protein phosphatase activity regulates Six1-Dach-Eya transcriptional effects in mammalian organogenesis. Nature 426, 247–254 (2003).

70. Go, M. J., Eastman, D. S. & Artavanis-Tsakonas, S. Cell proliferation control by Notch signaling in Drosophila development. Development 125, 2031–2040 (1998).

71. Dai, W. & Montell, D. J. Live imaging of border cell migration in drosophila. Methods Mol. Biol. 1407, 153–168 (2016).

72. Okajima, T., Xu, A. & Irvine, K. D. Modulation of notch-ligand binding by protein O-fucosyltransferase 1 and fringe. J. Biol. Chem. 278, 42340–42345 (2003).

